# GAD65Cre drives reporter expression in multiple taste cell types

**DOI:** 10.1101/2021.01.14.426727

**Authors:** Eric D. Larson, Aurelie Vandenbeuch, Catherine B. Anderson, Sue C. Kinnamon

## Abstract

In taste buds, Type I cells represent the majority of cells (50-60%) and primarily have a glial-like function in taste buds. However, recent studies suggest that they have additional sensory and signaling functions including amiloride-sensitive salt transduction, oxytocin modulation of taste, and substance P mediated GABA release. Nonetheless, the overall function of Type I cells in transduction and signaling remains unclear, primarily because of the lack of a reliable reporter for this cell type. GAD65 expression is specific to Type I taste cells and GAD65 has been used as a Cre driver to study Type I cells in salt taste transduction. To test the specificity of transgene-driven expression, we crossed GAD65Cre mice with floxed tdTomato and Channelrhodopsin (ChR2) lines and examined the progeny with immunochemistry, chorda tympani recording, and calcium imaging. We report that while many tdTomato+ taste cells express NTPDase2, a specific marker of Type I cells, we see expression of tdTomato in both Gustducin and SNAP25 positive taste cells. We also see ChR2 in cells just outside the fungiform taste buds. Chorda tympani recordings in the GAD65Cre/ChR2 mice show large responses to blue light, larger than any response to standard taste stimuli. Further, several isolated tdTomato positive taste cells responded to KCl depolarization with increases in intracellular calcium, indicating the presence of voltage-gated calcium channels. Taken together, these data suggest that GAD65Cre mice drive expression in multiple taste cell types and thus cannot be considered a reliable reporter of Type I cell function.

## INTRODUCTION

It is commonly assumed that taste buds consist of 3 cell types based on morphological, molecular and physiological characteristics(for review, see Roper and Chaudhari, 2017). Each cell type responds to different taste stimuli and communicates in various ways with the afferent nerve fibers to convey the message to the central nervous system. More specifically, Type II cells possess the receptors and downstream transduction components for sweet, umami and bitter stimuli. Although they do not form conventional synapses with nerve fibers, Type II cells release ATP via non-vesicular “channel” synapses to stimulate afferent gustatory nerve fibers(Romanov et al., 2018; Nomura et al., 2020). Conversely, Type III cells, which respond to acid stimuli and some salts, form conventional synapses with nerve fibers and release serotonin to activate neighboring cells or nerve fibers. Finally, Type I cells are the least understood cell type, representing ∼50% of the total number of cells in a taste bud (for review, see Roper and Chaudhari, 2017). Because they express NTPDase2 for the degradation of ATP released by other cells (Bartel et al., 2006; Vandenbeuch et al., 2013), and because they wrap around Type II and Type III cells (Yang et al., 2020), Type I cells are thought to have a glial-like function, similar to astrocytes in the nervous system. However, several studies have suggested a more complex function of Type I cells including a potential role in amiloride-sensitive salt taste transduction (Vandenbeuch et al., 2008, Baumer-Harrison et al., 2020), release of oxytocin to modulate taste (Sinclair et al., 2010), and release of GABA in response to trigeminal modulation with substance P (Huang and Wu, 2018). In order to examine the potential roles of Type I cells a specific marker is needed to both identify isolated cells for physiology and to manipulate gene expression.

GAD65 is expressed by mature Type I taste cells (Dvoryanchikov et al., 2011) and thus may be a good candidate to use as a Cre driver to report expression in Type I taste cells. GAD65Cre mice are available commercially and have been documented to drive expression specifically in GAD65 expressing neurons in the brain (Taniguchi et al., 2011). Taste cells, unlike neurons, turn over throughout life and since Cre is turned on at the birth of a cell and remains on, it is important to document its specificity of expression in adult taste cells. For example, if GAD65 is expressed transiently in taste cell progenitors, it may drive expression in more than just Type I taste cells. To examine more closely the specificity of expression, we obtained the commercially available GAD65Cre mice and crossed them with floxed Channelrhodopsin (ChR2) or Rosa tdTomato mice to examine specificity of expression. We used immunocytochemistry, chorda tympani nerve recordings, and calcium imaging of isolated tdTomato-positive taste cells. We conclude that while many Type I cells are labeled with either reporter, we show some expression of reporter in Type II and Type III taste cells, suggesting that GAD65 may be transiently expressed in some taste cell progenitors. Thus, this line cannot be used to drive reporter activity faithfully in Type I cells.

## MATERIALS AND METHODS

### Animals

All animal procedures in the study were approved by the Institutional Animal Care and Use Committee at the University of Colorado Anschutz Medical Campus. Genetically altered mice used in the study are described in Table 1. Male and female mice aged 4 − 8 months were used for this study. Mice were kept in passive air exchange caging under a 12h light/dark cycle and were provided food and water *ad libitum*. TRPM5GFP mice are described in (Clapp et al., 2006) and PKD2L1YFP mice are described in (Chang et al., 2010).

**Table 1.** Mouse lines used in this study. TRPM5GFP mice are described in (Clapp et al., 2006) and PKD2L1YFP mice are described in (Chang et al., 2010).

| Name | Jax stock | Jax full name | background |
| --- | --- | --- | --- |
| GAD65Cre | 010802 | Gad2-IRES-Cre | C57BL/6J |
| tdTomato | 007909 | Ai9(RCL-tdT) | C57BL/6J |
| ChR2 | 024109 | Ai32(RCL-ChR2(H134R)/EYFP) | C57BL/6J |
| TRPM5GFP | n/a | n/a | C57BL/6J |
| PKD2L1YFP | n/a | n/a | C57BL/6J |

### Immunohistochemistry

Tissues were fixed by immersion (4h at 4C) or transcardial perfusion followed by post-fixation (4h at 4°C) with 4% paraformaldehyde dissolved in 0.1M Phosphate Buffer (PB, pH 7.2). Fixed tissues were cryoprotected overnight at 4°C in 20% sucrose dissolved in 0.1M PB prior to embedding in Optimal Cutting Temperature Compound (Tissue-Tek, Torrance, CA). Twelve-16 µm sections were cut using a cryostat (Leica) and collected on charged glass microscope slides (SuperFrost Plus, Fisher Scientific). Slides were stored at −20°C until use. After thawing, slides were rinsed with PBS and incubated in blocking solution (2% normal donkey serum, 1% bovine serum albumin, 0.3% triton in PBS) prior to overnight incubation in primary antibody solution. Antisera used in this study are included in Table 2 and were diluted in blocking solution. Secondary antisera were diluted in blocking solution and applied for 3h at room temperature. Slides were counterstained with DAPI (0.5 ug/mL, ThermoFisher) and coverslipped with FluoroMount-G (Southern Biotech).

**Table 2.**
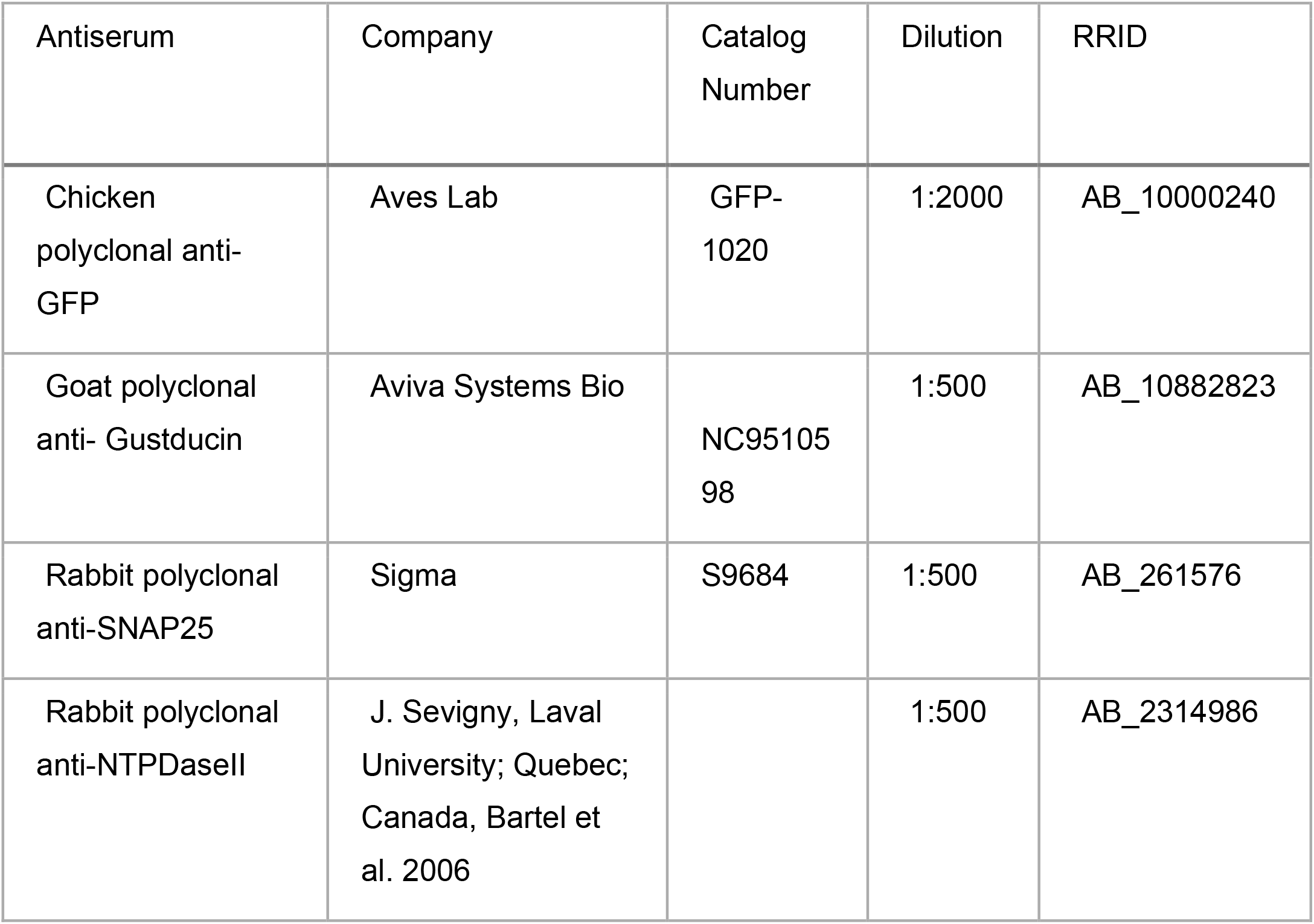
List of primary antisera used in this study.

#### Whole mount IHC

To image whole mount fungiform taste buds, fungiform epithelia were peeled as described below, pinned flat and fixed in 4% paraformaldehyde for 1 hour at room temperature. IHC protocol was like that of tissue sections, but with increased timing: overnight blocking at 4C, 48h primary antibody at 4C, overnight secondary antibody at 4C. Tissue was mounted on a microscope slide using FluoroMount-G.

#### Confocal Imaging

All images were acquired using a Leica SP8 laser scanning confocal microscope with a 63x oil immersion objective (NA1.4). Images were acquired with 1024×1024 pixel resolution and axial intervals of ∼0.3 µm. Fluorescent signals were detected sequentially by line to limit any cross-channel detection except for DAPI and Alexa-647 which were acquired simultaneously. Images were processed using a combination of ImageJ (FIJI SCR_002285) and the Adobe CC platform (San Jose, CA). Manipulation of images included only linear modifications to the minimum and maximum pixel values, changes to pseudocolor, cropping, and pixel density resampling ([Photoshop:Image Size: resample] to achieve appropriate resolution for display).

#### Image analysis

Cell profile intensities were measured by creating ROIs around cell profiles at multiple positions through the Z-stack, with ∼5-7 um axial spacing between positions (and excluding the first and last 5 um of whole-mount images) using ImageJ. The mean intensity per ROI was calculated for each channel at each axial position. For each image, background values and maximum values were calculated using a maximum z-projection of optical planes used in the analysis above. For background values, 5 ROIs were drawn in areas directly adjacent to the taste bud, and the mean pixel intensity values were measured. Threshold for each channel was calculated as mean background plus 2 standard deviations. Maximum values were calculated by determining the pixel intensity value that represented 95.5% of non-zero pixels, per channel. Measurements from ImageJ were imported to R (R Core Team, 2019) using a custom script where background and cell profile levels were scaled based on the maximum pixel values, summarized, and plotted using *ggplot2* (Wickham, 2016).

### Taste Cell Isolation

Circumvallate and fungiform taste buds and cells were isolated as previously described (Béhé et al., 1990; Clapp et al., 2006). Mice were euthanized with CO_2_ followed by cervical dislocation. An enzyme cocktail consisting of 1 mg/ml collagenase A (Roche, Indianapolis, IN), 3 mg/ml Dispase II (Roche, Indianapolis, IN), and 1 mg/ml trypsin inhibitor (Sigma, St. Louis, MO) dissolved in Tyrode’s (140 NaCl, 5 KCl, 1 MgCl2, 1 CaCl2, 10 HEPES, 10 glucose, and 1 pyruvate (pH 7.4 with NaOH)) was injected underneath the epithelium of the tongue. After incubation for 40 min. in oxygenated Ca-Mg free Tyrode’s (same as above but 1 mM BAPTA replaced the divalent cations), the epithelium was separated from the underlying connective tissue and placed in Ca-Mg free Tyrode’s for 30 min. Taste buds were removed by gentle suction with a fire-polished pipette and plated onto glass coverslips coated with Cell Tak (BD Biosciences, Bedford, MA) or poly-L-lysine (Sigma, St. Louis, MO). Coverslips were placed in a perfusion chamber (RC-25F, Warner Scientific Inc., Hamden, CT). All data for calcium imaging were obtained from isolated taste cells or small cell clusters to verify the presence or absence of tdTomato expression.

### Calcium imaging

10 minutes after plating on Cell-Tak coated coverslips, cells were loaded with 2 μM fura-2 AM (Invitrogen, Waltham, MA) for 25 minutes at 35°C. After loading, cells were washed and kept in a continuous flow of Tyrode’s using a gravity flow perfusion system (Automate Scientific Inc., San Francisco, CA). Depolarizing concentrations of KCl (55 mM) were made in Tyrode’s (with 90 mM NaCl to maintain osmolarity). Sequential excitation at 340 and 380 nm from an Hg light source was controlled using a Sutter Lambda 10-3. Emission at ∼ 510 nm was measured using a Retiga R3 (QImaging) and a 40x oil immersion objective (NA1.4) on an inverted Olympus IX71. Images were captured every 3 seconds using Imaging Workbench 6.1 (Indec Biosystems,Inc). Baseline ratio per cell was measured as the mean ratio 20 seconds pre- and post-stimulus application. In the event a cell did not return to baseline, the first 20 seconds of a stable ratio was used as the post-stimulus baseline. If the post-stimulus baseline was greater than 25% of the response amplitude, the cell was removed from analysis. A ratio of emission at 510 from excitations at 340 and 380 nm was considered a response if it fell 2 standard deviations or more above the median baseline ratio across all cells.

### Nerve recording

Mice were anesthetized with urethane (2 g/kg; i.p), maintained in a head holder, and a cannula was placed in the trachea to facilitate breathing. The chorda tympani nerve was exposed using a ventral approach, cut near the tympanic bulla and placed on a platinum-iridium wire. A reference electrode was placed in a nearby tissue. The signal was fed to an amplifier (P511; Grass Instruments), integrated, and recorded using AcqKnowledge software (Biopac). The anterior part of the tongue was stimulated with different tastants (applied for 30 s and then rinsed with water for 40 s) with a constant flow pump (Fisher Scientific). Stimuli consisted of (in mM): NH_4_Cl 100, quinine 10, sucrose 500, NaCl 100, citric acid 10. For light stimulation, a LED blue light (470nm) was positioned in close proximity to the tongue. Light pulses were chosen to give an optimal nerve response (5Hz, 7mW, 50% duty cycle; Wilson et al., 2019; Vandenbeuch et al., 2020). The amplitude of each integrated response was averaged over the 30 s application using AcqKnowledge software and subtracted to the average baseline activity averaged 5 s before tastant application. In order to average responses from different animals, each response was normalized to baseline using the following formula: (amplitude response - baseline) / baseline)(Vandenbeuch et al., 2015; Larson et al., 2020). A Grubb’s test was used to eliminate significant outliers (GraphPad). Responses were compared using a one-way ANOVA with Tukey’s post hoc test.

## RESULTS

### Immunohistochemical characterization of GAD65Cre

To use a genetic model to study Type I cell function, the Cre driver allele of interest *must* be specific to Type I cells in adult mice. Thus, we carefully characterized the specificity of Cre recombinase in GAD65Cre mice using floxed tdTomato and ChR2 reporter alleles. We additionally crossed GAD65Cre/tdTomato with TRPM5GFP and PKD2L1YFP (to label most Type II and III cells with GFP/YFP (Clapp et al., 2006; Chang et al., 2010; Wilson et al., 2017)). GAD65Cre-driven tdTomato and ChR2 were robustly expressed in taste buds of multiple taste fields (including fungiform and CVP (figure 1-4) and foliate and soft palate (not shown). To determine if labeled cells were Type I cells, we labeled lingual sections from GAD65Cre/tdTomato mice with antibodies against NTPDase2, a Type I cell marker (Bartel et al., 2006). NTPDase2-immunoreactivity localizes to Type I cell membranes and often coincided with the tomato reporter, suggesting that tomato+ cells are indeed Type I cells (Fig 1). NTPDase2 immunoreactivity also often colocalized with ChR2 (not shown), which is a membrane-associated reporter, though colocalization of NTPDase2 and ChR2 does not necessarily mean they are in the same cell; signals from two adjacent (abutting) cell membranes would result in artefactual colocalization given resolution limits of light microscopy (Corson and Erisir, 2013; Stratford et al., 2017). Thus, we chose not to pursue quantitative analysis here. Qualitatively, most, but not all, NTPDase2 immunoreactive cells were associated with either tdTomato or ChR2 fluorescence. Conversely, not all tdTomato or ChR2 signals were associated with NTPDase2 immunoreactivity. This suggested that another, unique cell type is labeled with the reporter or that the reporter may be expressed in a subset of Type II and/or Type III cells. Additionally, ChR2+ cells were observed around the circumference of some but not all fungiform taste buds (5/8 taste buds with ChR2, Figure 2). These cells were not immunoreactive to any other markers used in this study. Interestingly, these cells were observed less frequently (and less brightly) when tdTomato was used as the reporter (not shown). Stimulation of ChR2 in these cells has the potential for unforeseen effects. Taken together, these data support that Cre recombinase is expressed in Type I cells but is not limited to Type I cells.

**Figure 1.**
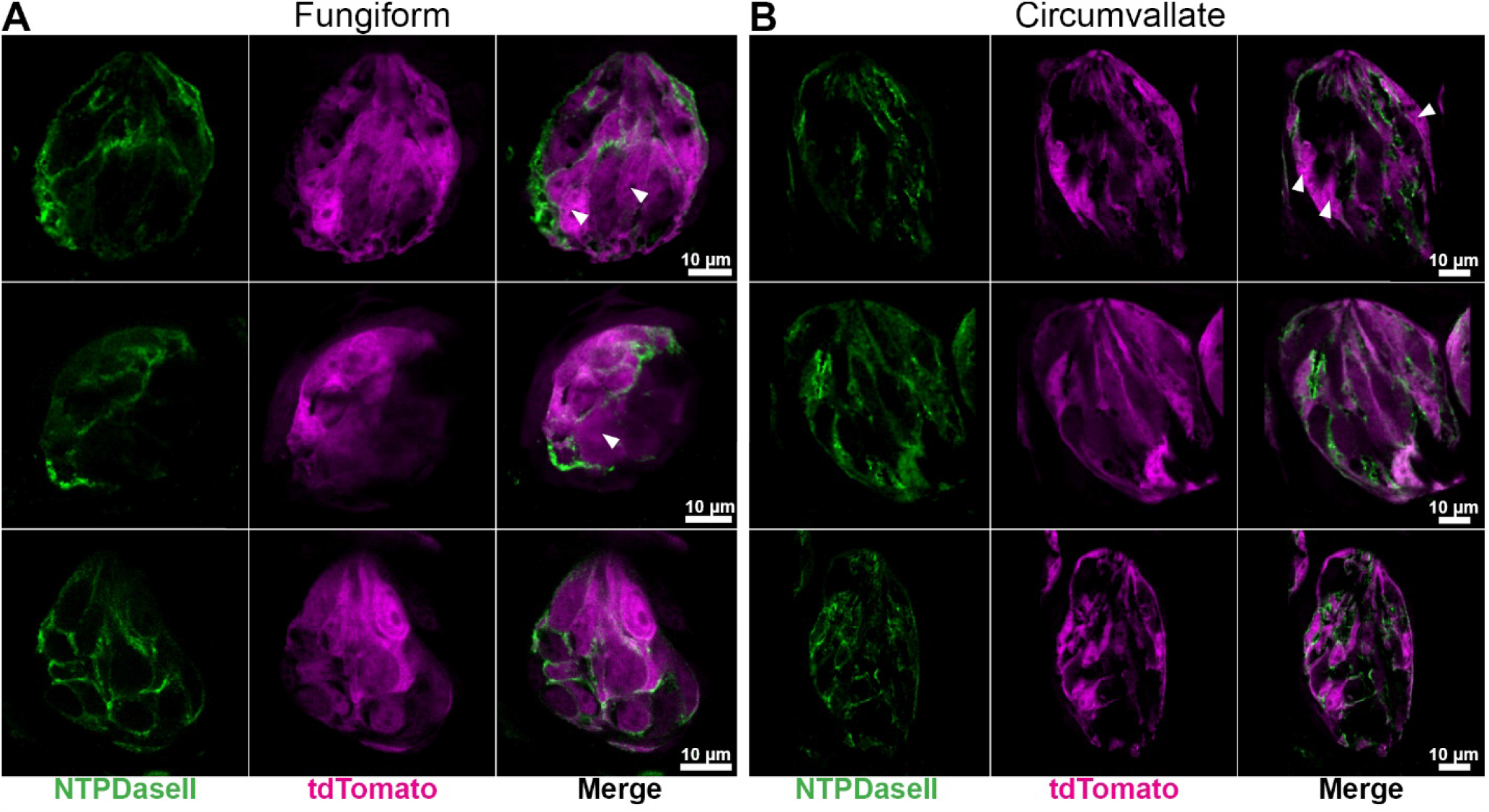
2 column figure, width 7.09 inches Confocal images of fungiform (A) and circumvallate (B) tastebuds from GAD65Cre/tdTomato mice. Images are single optical planes. Six representative taste buds sectioned longitudinally show NTPDase2 immunoreactivity (green) and endogenous tdTomato fluorescence (magenta) in the taste bud. NTPDase2 immunoreactivity is mostly associated with tdTomato fluorescence. Arrowheads show areas of tdTomato not associated with strong membrane-localized NTPDase2 immunoreactivity. Scale bars, 10 μm.

**Figure 2.**
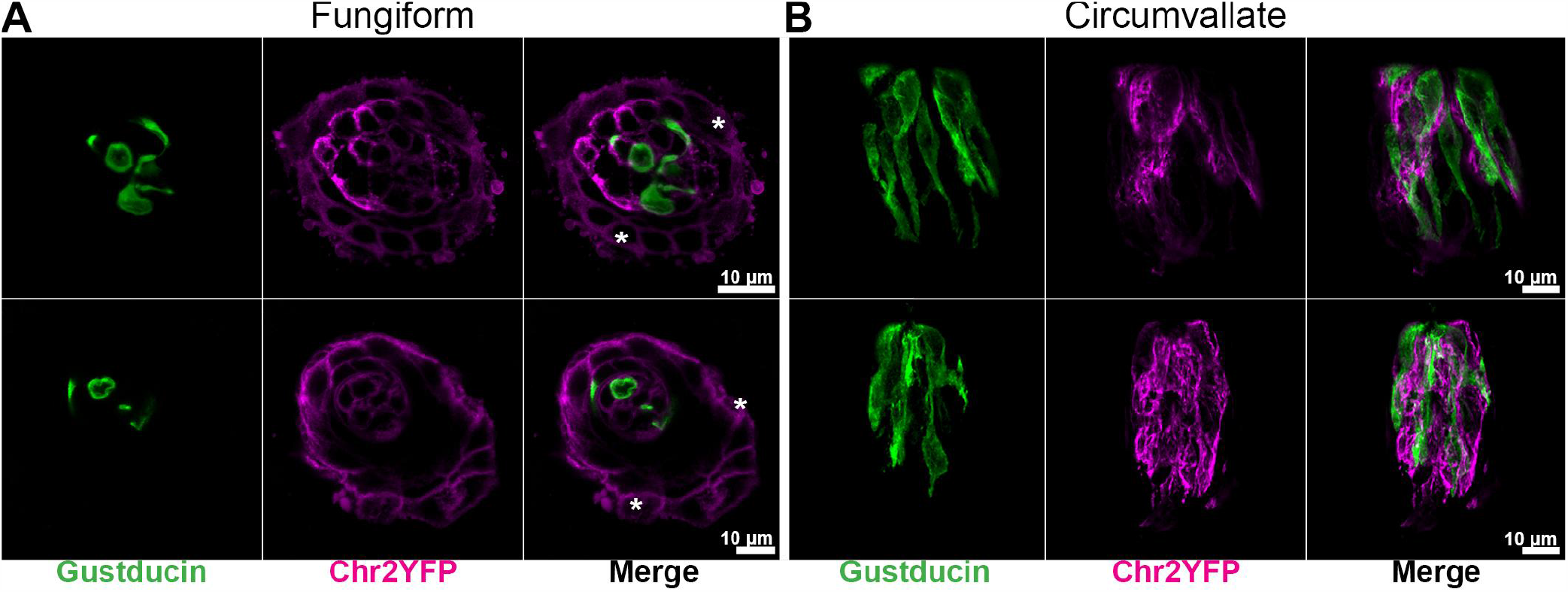
2 column figure, width 7.09 inches Confocal images of whole-mount fungiform (A) and longitudinally sliced circumvallate (B) taste buds from GAD65Cre/ChR2. Fungiform images are single optical planes while CV images are a Z-stacks of 2 axial planes spaced 0.75 μm apart. In many GAD65Cre/ChR2 fungiform taste buds, a ring of ChR2+ cells was observed outside the taste bud (asterisks). The identity of these cells is unclear. Additionally, while ChR2 was observed surrounding Gustducin+ cells, it is unclear if the signals are in the same cell, given the proximity of cells with one another in taste buds.

To examine the taste cell type-specificity of GAD65Cre activity further, we examined co-expression of tdTomato with Type II and Type III cell markers, Gustducin and SNAP25, respectively. In 7 whole-mount fungiform taste buds from GAD65Cre/tdTomato mice, 4 had at least 1 cell where above-threshold tdTomato fluorescence was detected in a Gustducin-immunoreactive cell, and 4 had at least 1 cell where above-threshold tdTomato fluorescence was detected in a SNAP25 immunoreactive cell (Figure 3). Quantitative analysis of whole mount fungiform epithelia revealed that of 498 cell profiles analyzed, 20 (4.0%) had above threshold levels of both tdTomato fluorescence and Gustducin immunoreactivity, and 13 (2.6%) had above threshold levels of tdTomato fluorescence and SNAP25 immunoreactivity. The remaining ROIs included 183 (36.7%) tdTomato+, 170 (34.1%) Gustducin+, 46 (9.2%) SNAP25+, and 66 (13.3%) sub-threshold profiles. SNAP25-immunoreactivity and Gustducin-immunoreactivity were never observed in the same profile (not shown). These data suggest that a small proportion of type II and III cells are also labeled by GAD65Cre.

**Figure 3.**
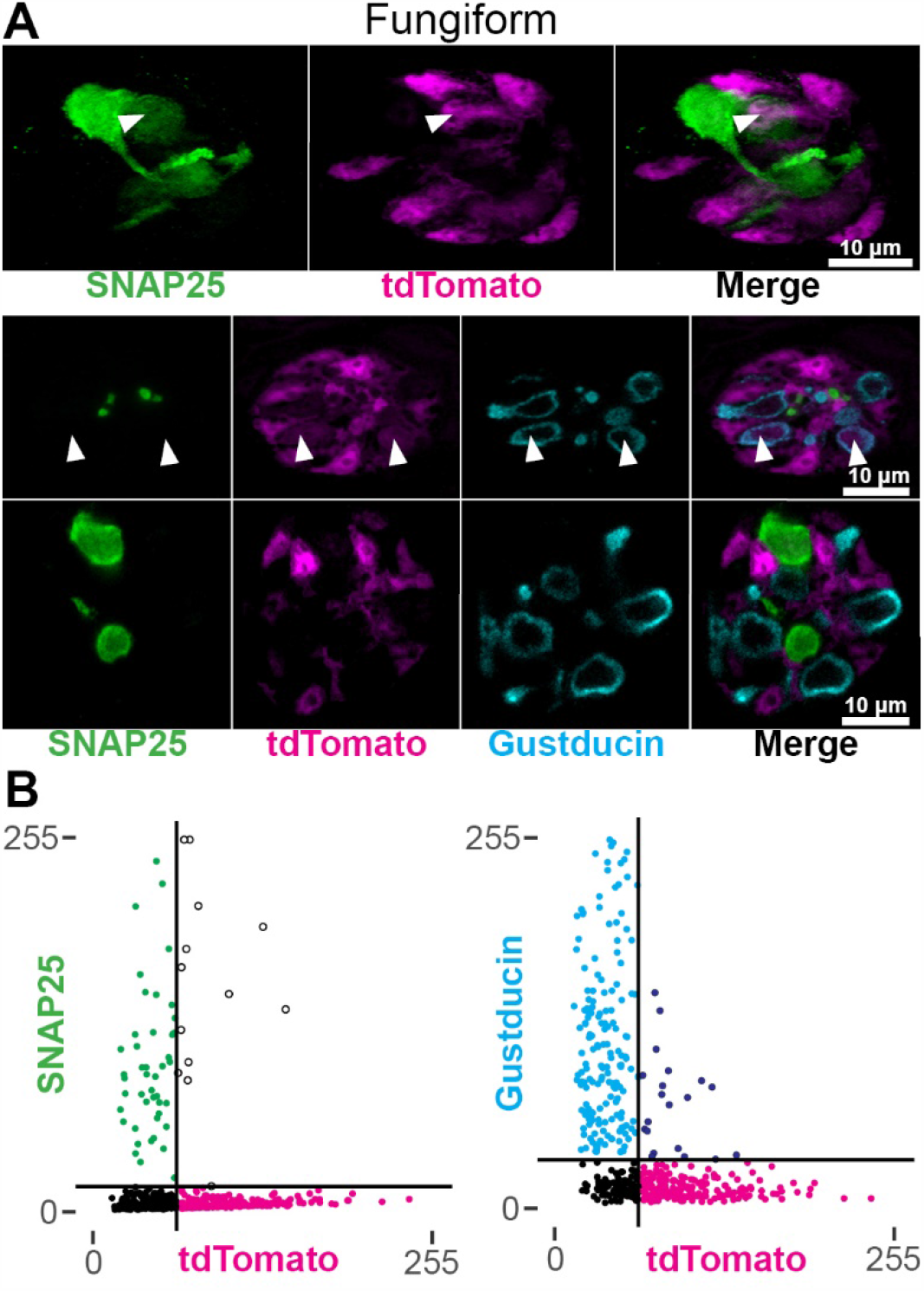
1 column figure, width 3.54 inches A. Confocal images of whole-mount fungiform taste buds from GAD65Cre/tdTomato mice. Fungiform taste buds are imaged in cross section. Images are single optical planes. Type III cells are identified by SNAP25 immunoreactivity (green) and a subset of type II cells are identified by Gustducin immunoreactivity (cyan). Arrowheads indicate tdTomato fluorescence in either a Type II or Type III cell. Images were background-subtracted and scaled using the same method for quantification (see methods). B. Quantification of cell profile fluorescence intensities. Cell profiles were measured in optical sections to assess the degree of overlapping signals. Horizontal and vertical black lines show the calculated threshold levels of 2 standard deviations above background. Black points are those below threshold in both channels. Points in the upper right quadrant are those found above-threshold in both channels.

To eliminate any confounds of antibody-labeling and to maximize the number of Type II and III cells identified, we examined taste buds from GAD65Cre/tdTomato/TRPM5GFP/PKD2L1YFP mice. GFP and YFP fluorophores in these tissues were detected using an anti-GFP antibody (cross-reacts with YFP), allowing us to visualize (but not distinguish) Type II and III cells in the same channel. A subset of Type II cells was further identified by Gustducin immunoreactivity. In 8 whole mount fungiform taste buds that were imaged, 6 taste buds had at least one Type II or III cell that coincided with tdTomato (Fig 4). These observations were consistent with antibody labeling in GAD65Cre/tdTomato mice and held true for other taste fields including posterior regions (circumvallate and foliate papilla). We observed tdTomato expression in 13.5% of fungiform, 3.7% of soft palate, 5.4% of foliate, and 5.8% of circumvallate cells immunoreactive for TRPM5GFP/PKD2L1YFP and/or Gustducin (Table 3).

**Figure 4.**
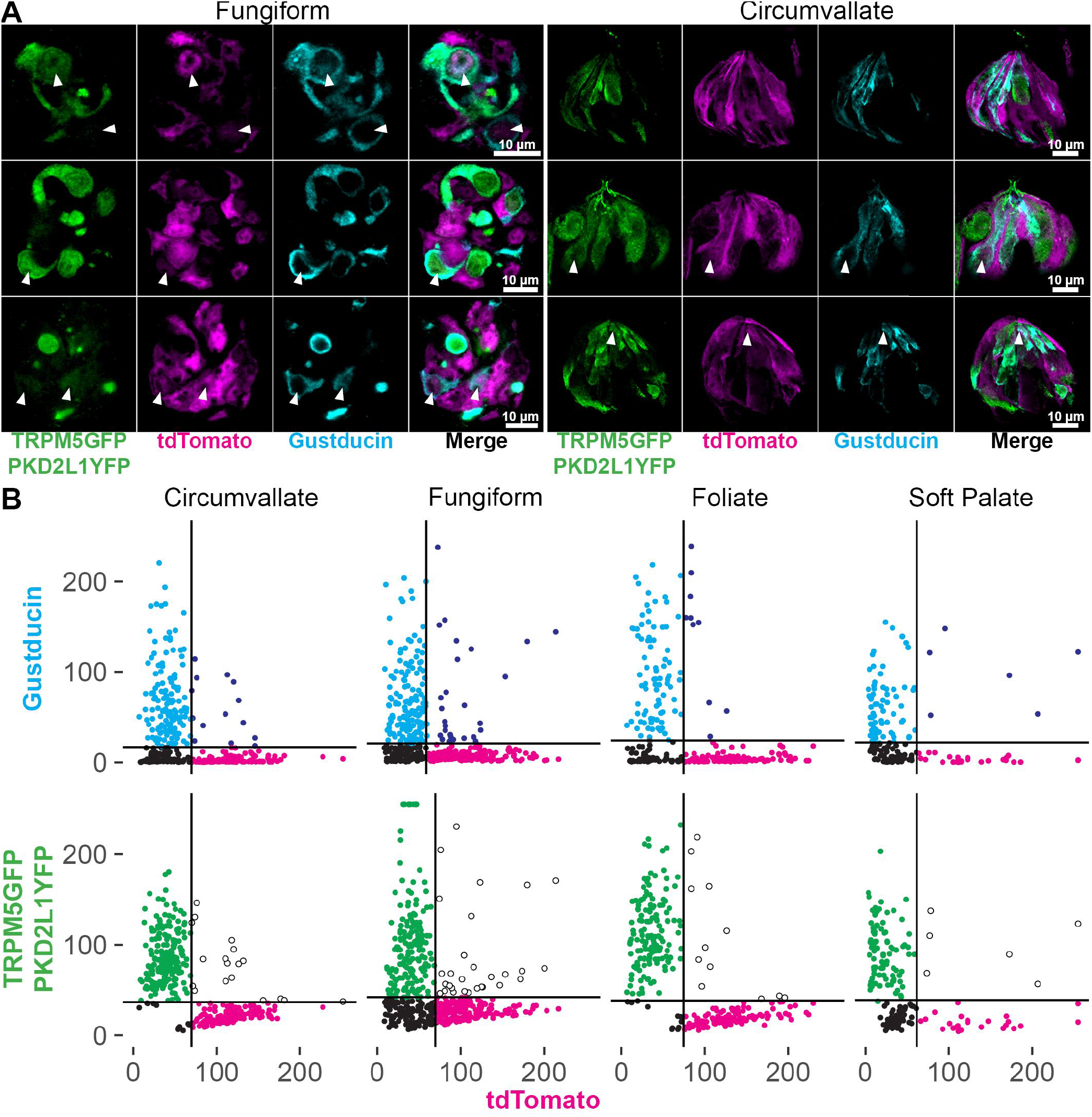
2 column figure, width 7.09 inches Confocal images of whole-mount fungiform taste buds from GAD65Cre/tdTomato/TRPM5GFP/PKD2L1YFP mice. Taste buds are imaged in cross section. Images are single optical planes. Type II and III cells are visualized with GFP immunoreactivity as the antibody reacts with both GFP and YFP. A subset of type II cells is visualized by Gustducin immunoreactivity. tdTomato fluorescence in a Type II/III cell is observed in at least 1 cell of each taste bud (arrowheads). B. Quantification of cell profile mean fluorescence intensities. Cell profiles were measured in optical sections to assess the degree of overlapping signals. Horizontal and vertical black lines show the calculated threshold levels of 2 standard deviations above background. Black points are those below threshold in both channels. Points in the upper right quadrant are those found above-threshold in both channels.

**Table 3.**
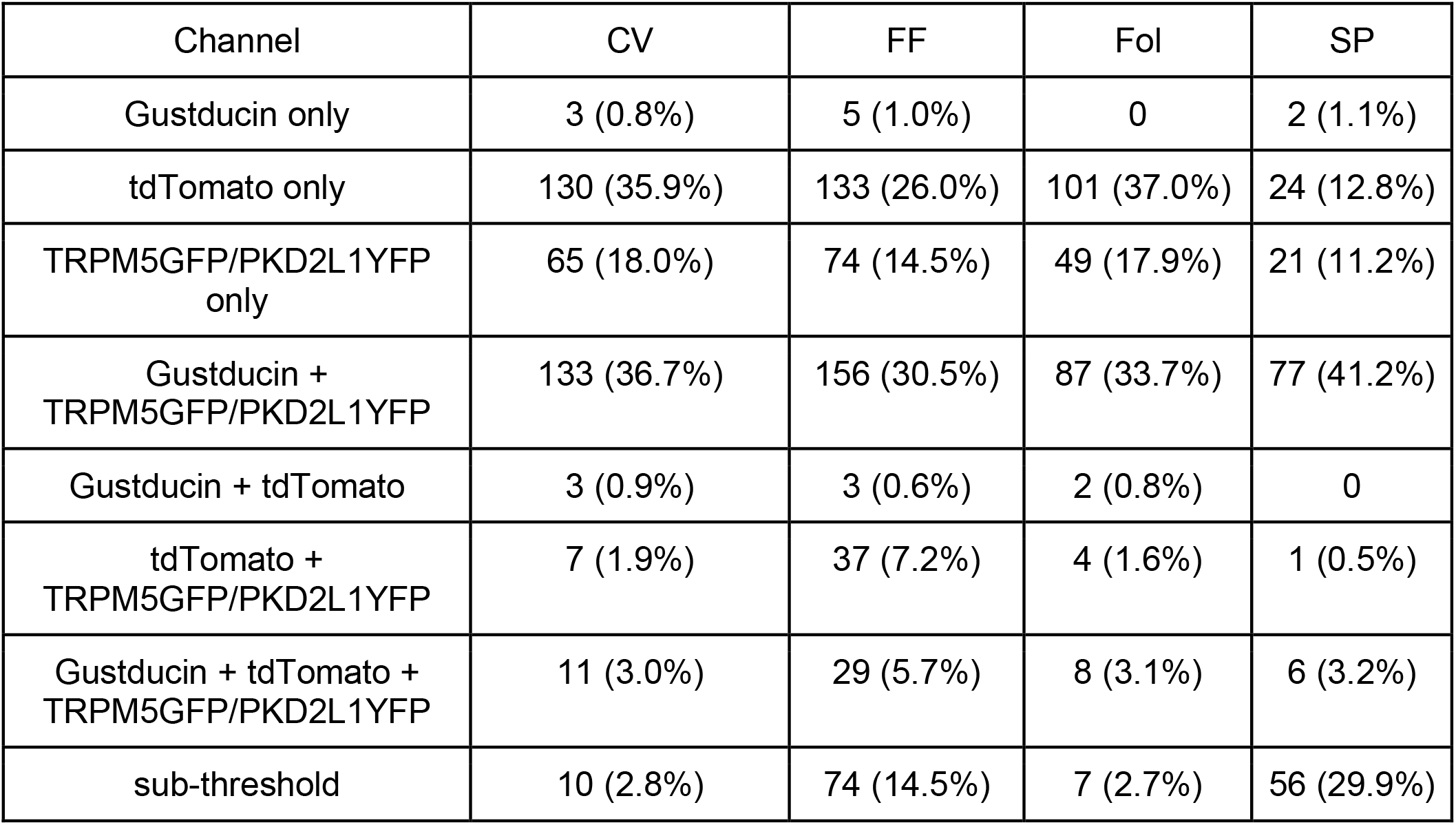
Summary of quantification in Figure 4. Counts are number of ROIs displaying above-threshold pixel values for each channel or channel combination. Percentages are reflective of percent ROIs from each taste field. Each cell profile is binned according to channel, thus a single cell profile is counted only once.

| Channel | CV | FF | Fol | SP |
| --- | --- | --- | --- | --- |
| Gustducin only | 3 (0.8%) | 5 (1.0%) | 0 | 2 (1.1%) |
| tdTomato only | 130 (35.9%) | 133 (26.0%) | 101 (37.0%) | 24 (12.8%) |
| TRPM5GFP/PKD2L1YFP only | 65 (18.0%) | 74 (14.5%) | 49 (17.9%) | 21 (11.2%) |
| Gustducin + TRPM5GFP/PKD2L1YFP | 133 (36.7%) | 156 (30.5%) | 87 (33.7%) | 77 (41.2%) |
| Gustducin + tdTomato | 3 (0.9%) | 3 (0.6%) | 2 (0.8%) | 0 |
| tdTomato + TRPM5GFP/PKD2L1YFP | 7 (1.9%) | 37 (7.2%) | 4 (1.6%) | 1 (0.5%) |
| Gustducin + tdTomato + TRPM5GFP/PKD2L1YFP | 11 (3.0%) | 29 (5.7%) | 8 (3.1%) | 6 (3.2%) |
| sub-threshold | 10 (2.8%) | 74 (14.5%) | 7 (2.7%) | 56 (29.9%) |

Overall, we hypothesize that the expression of Cre recombinase in more than just Type I cells in these mice could be the result of a shared stem/precursor population of adult taste cells (Castillo et al., 2014; Gaillard et al., 2015). Thus, activation of Cre recombinase could result in the constitutive reporter expression in adult taste cells that no longer express GAD65, as well as in Type I taste cells known to express mRNA coding for GAD65 (Dvoryanchikov et al. 2011).

### Calcium imaging of isolated GAD65+ cells

Type III cells are the only cells in the taste bud with voltage-gated calcium channels (Medler et al., 2003; Romanov and Kolesnikov, 2006; Vandenbeuch et al., 2010), hence isolated Type III cells are identifiable by robust calcium influx in response to depolarization with KCl. As expected, a subset of tdTomato negative cells exhibited calcium responses to KCl (CV:5/15, FF: 3/7). However, a subset of tdTomato+ cells also had moderate to robust calcium responses to KCl (CV:7/19, FF: 3/13), consistent with reporter expression in Type III taste cells (Fig. 4).Thus, these physiological data further support the argument that GAD65Cre does not exclusively label Type I cells.

### Chorda tympani responses elicited by blue light

When blue light is applied to the anterior tongue, robust chorda tympani responses are observed in GAD65Cre/ChR2 mice (Fig. 5). The amplitude of the responses to blue light (average ratio to baseline ± sem: 3.07 ± 0.41) are larger than any other individual tastant used including NH_4_Cl 100mM (1.29 ± 0.16), sucrose (0.48 ± 0.07), NaCl (0.99 ± 0.33), citric acid (0.45 ± 0.03), and quinine (0.33 ± 0.41) applied to the same region of the tongue. Responses to blue light are significantly different from all other tastants (One way ANOVA with Tukey’s post-hoc; F(1, 5) = 20.06, p < 0.001). The large light responses are likely due to the large number of cells stimulated (including Type II and III cells). These data further support our contention that GAD65Cre drives ChR2 expression in multiple taste cell types.

**Figure 5.**
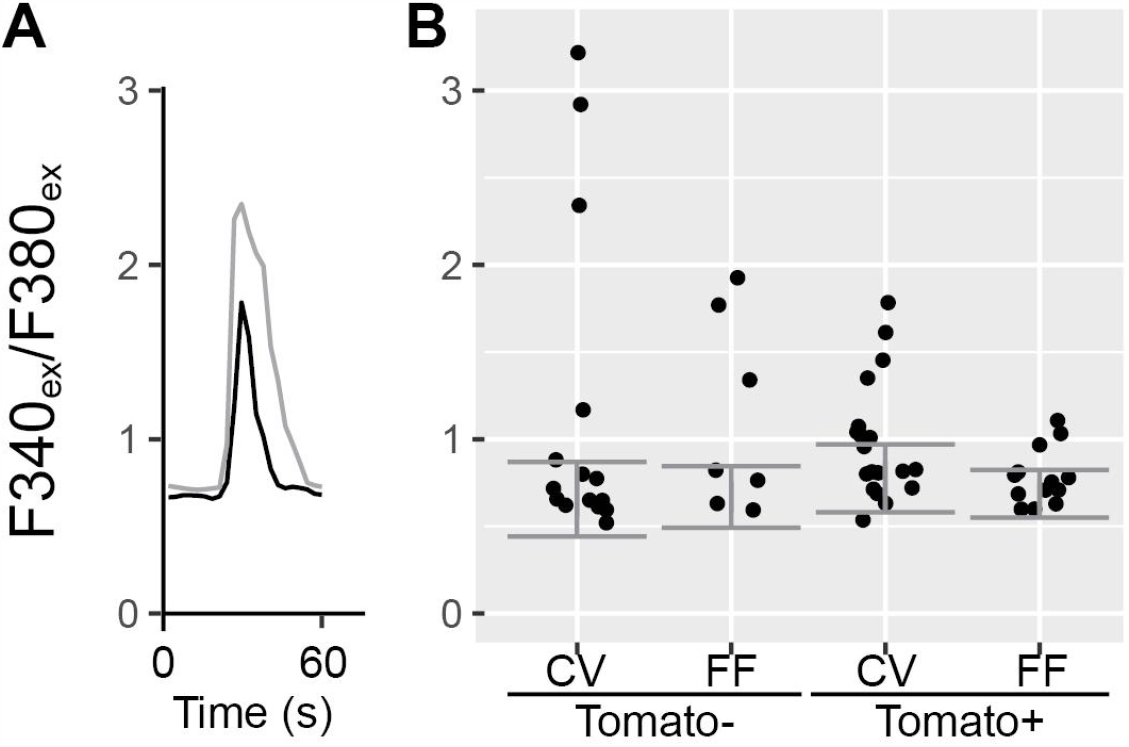
1 column figure, width 3.54 inches Fura2 calcium imaging of isolated GAD65Cre/tdTomato cells. tdTomato positive and negative taste cells were loaded with Fura2 and exposed to depolarizing concentrations of KCl to test for the presence of voltage gated calcium channels. Responses are displayed as a ratio of emission at 510 from excitations at 340 and 380 nm. A. Two example cells showing a response to KCl (black: tdTomato positive, grey: tdTomato negative). B. Responses of individual cells. Horizontal position of each dot within each group is arbitrary: dots are jittered to visualize overlapping points. Horizontal error bars depict 2 standard deviations above and below median of baseline values across all cells per group. Responses over median plus 2 standard deviations were considered positive.

**Figure 6.**
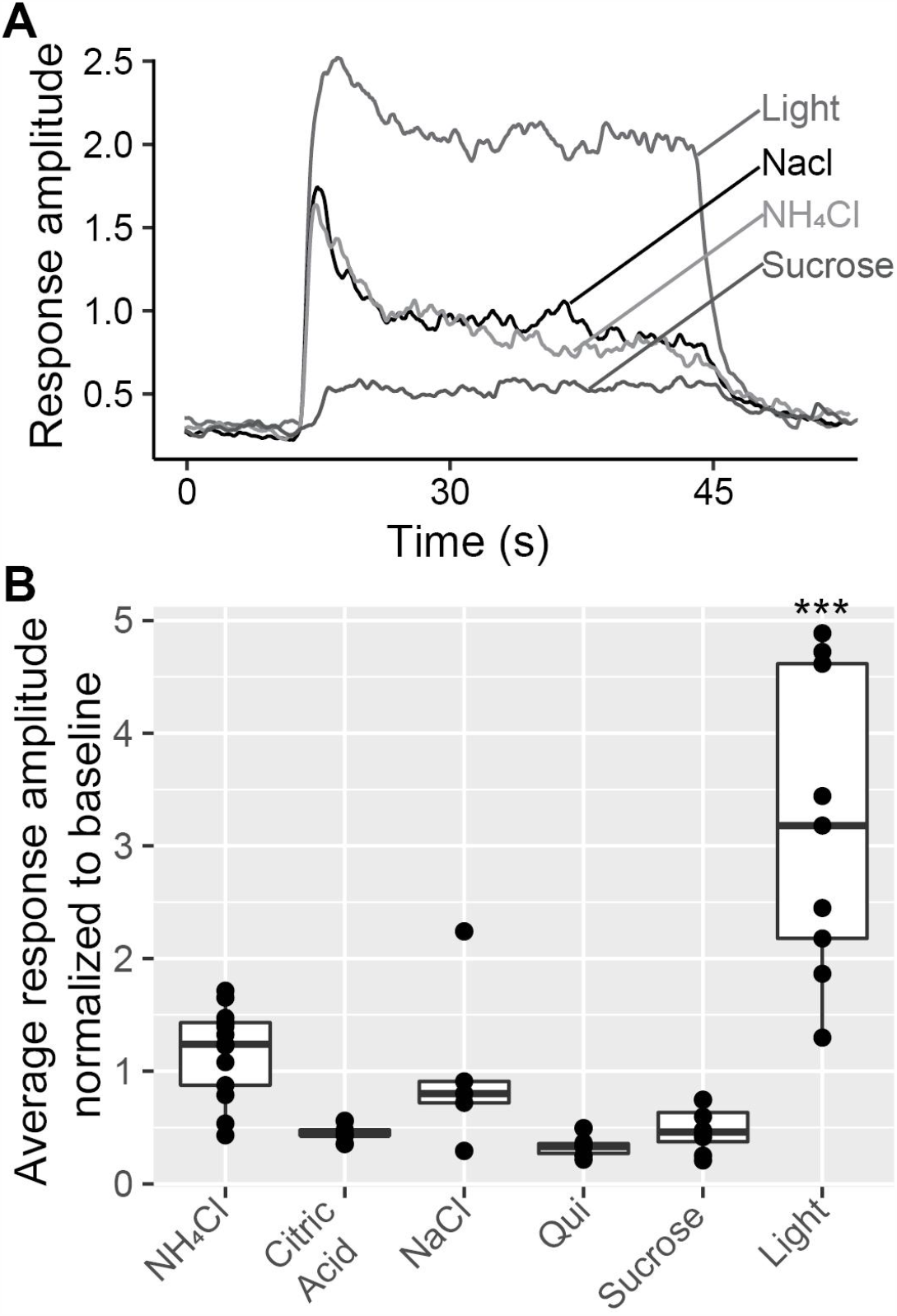
1 column figure, width 3.54 inches Chorda tympani nerve recordings in GAD65Cre/ChR2 mice. A: Representative integrated nerve recordings in response to light (5Hz), NaCl 100mM, NH_4_Cl 100mM, and sucrose 500mM. B: Average chorda tympani responses normalized to baseline. Dots represent individual animals and the box plot demonstrates distribution of data. *** indicates significance at p<0.001 compared to all other tastants by One way ANOVA with Tukey’s post-hoc. No differences among tastants were observed.

## DISCUSSION

The main goal of the present study was to characterize a commercially available mouse line that expresses Cre recombinase from the *Gad2* promoter (GAD65 protein), to evaluate use of this allele to mark and/or manipulate gene expression in Type I taste cells. GAD65, encoded by the *Gad2* gene, is a Glutamate decarboxylase 2 enzyme which catalyzes production of gamma-aminobutyric acid (GABA) from L-glutamic acid. These mice have been used to examine the role of GABA and GAD65 in the brain where GAD65Cre-driven reporter expression was specific for GABA-expressing neurons with no mis-expression in other neuron types (Dergacheva et al., 2020; Quina et al., 2020; Whyland et al., 2020). This specific labeling is consistent with the observation that GABAergic neurons turn on *Gad2* as they differentiate (Westmoreland et al., 2001). However, GAD65Cre activity has not been fully characterized in the peripheral taste system, where taste cells are continuously generated throughout adult life (for a review, see (Barlow, 2015)). Here, we crossed GAD65Cre mice with floxed tdTomato or channelrhodopsin (ChR2) mice to identify and/or stimulate lineage traced cells. Although a majority of tdTomato or ChR2 expressing cells also expressed the Type I cell-marker, NTPDase2, immunohistochemical analyses demonstrate expression of the GAD65Cre-driven reporter in subsets of both Type II and Type III cell types as well. This misexpression may be due to expression of Cre recombinase in taste cell progenitors, if such progenitor cells transiently express GAD65. Thus, if Cre recombinase is turned on in these progenitors, the progeny will continue to express the reporter, even though adult taste buds only express GAD65 in Type I cells.

Physiological studies also suggest that reporter expression is not limited to Type I taste cells. First, calcium imaging experiments showed that a significant subset of isolated GAD65Cre/tdTomato cells responded to KCl depolarization with increases in intracellular calcium. The only adult taste cells known to exhibit voltage-gated calcium channels are the Type III taste cells that utilize calcium dependent vesicular release of transmitter to activate sensory afferent nerve fibers (Medler et al., 2003; DeFazio et al., 2006; Roberts et al., 2009; Vandenbeuch et al., 2010). Although Dvoryanchikov et al (2011) reported *Gad2* mRNA (coding for GAD65 protein) expression in Type III adult taste cells (1 /18 cells in CV) by single cell PCR, our data show a much larger subset of isolated tdTomato cells with depolarization-evoked calcium increases (7/19 in CV (Fisher’s Exact two sided P value = 0.042 vs 1/18 cells from Dvoryanchikov et al); 3/13 in fungiform (not compared - Dvoryanchikov et al. did not examine fungiform taste cells), suggesting that GAD65Cre likely drives reporter expression in some Type III cell progenitors. Second, our chorda tympani nerve recordings show very large responses when blue light is applied to the tongue -- responses larger than any evoked by a taste stimulus. Although this does not rule out Type I cell stimulation activating nerve fibers, it does suggest that ChR2 is driven in multiple cell types that all are activated by light.

In a recent study, (Baumer-Harrison et al., 2020) used the same constitutive GAD65Cre mice crossed with floxed ChR2 mice to optogenetically stimulate taste buds. To examine reporter expression in taste buds, they used immunocytochemistry to label Type II (PLCb2) and Type III cells (Car4), and counted cells to determine co-localization with the reporter. No localization of ChR2 was observed in any cells that labeled with either Car4 or PLCb2, suggesting specificity of the reporter to Type I cells. Clearly these data differ from our results using the same Cre driver. We used different markers for Type II cells (GNAT vs PLCb2) and Type III cells (SNAP25 vs Car4), a different reporter for counting cells (tdTomato vs ChR2), and importantly, high-resolution confocal microscopy, and cross-sectional imaging of fungiform taste buds. We chose the tdTomato reporter for our studies as it is challenging to assess the degree of co-expression when comparing membrane-localized ChR2 to cytoplasmic immunoreactivity for Type II and III markers. With the large number of cells expressing ChR2, we were unable to say with certainty where individual cell boundaries began and ended. Thus, we used overlap of cytoplasmic tomato fluorescence with GFP and YFP expressed cytosolically in type II and III cells to determine with certainty whether GAD65Cre results in labeling of subsets of type II and III taste cells.

With regards to our cell profile counts, we may be underestimating the number of Type II or Type III cells that express tdTomato. For instance, in taste buds labeled with Gustducin (Figure 3) we are likely underestimating the number of Type II cells, as gustducin only identifies a subset of Type II cells (Clapp et al., 2001). Thus, some cells identified as tdTomato+ only could in fact be Gustducin-negative Type II cells that also express tdTomato. In taste buds of GAD65Cre/tdTomato/TRPM5GFP/PKD2L1YFP mice, while we are likely identifying nearly all Type II cells with TRPM5GFP, we may be underestimating the number of Type III cells. While PKD2L1YFP reliably identifies the majority of Type III cells in all taste fields (Wilson et al., 2017), the YFP we are detecting is tethered to the PKD2L1 protein which is predominantly localized in apical regions of Type III cells. Cell profiles were counted in 2-3 planes of the taste bud, spaced ∼10 um apart. Thus, in more basal planes, PDK2L1YFP is not a reliable identifier of Type III cells. Further, overall, we could be underrepresenting the cell counts given the high thresholds we are using for expression, especially the fungiform taste buds where ∼14% of profiles analyzed were below threshold in every channel. Thus, we predict that the GAD65cre is expressed in a higher percentage of Type II or Type III cells than calculated here.

Baumer-Harrison et al also reported that blue light evoked responses in sodium-best neurons of the nucleus of the solitary tract and drove salt appetitive behavior. They concluded that amiloride-sensitive salt responses were mediated by Type I taste cells. Our data would argue that while Type I cells were likely stimulated in their experiments, other cell types were also stimulated and it is not clear which cells were actually responsible for amiloride-sensitive salt taste. Another recent study also sheds doubt on Type I cells mediating amiloride-sensitive salt taste. (Nomura et al., 2020) showed that while amiloride-sensitive currents were present in a subset of Type I taste cells, validated by expression of NTPDase2, they did not signal to the nervous system. Instead, a subset of cells co-expressing the amiloride-sensitive Na channel, the ATP release channel CALHM1, and voltage-gated sodium channels were required for amiloride-sensitive salt taste chorda tympani responses. Type I cells express neither Calhm1 (Taruno et al., 2013) nor voltage-gated sodium channels (Medler et al., 2003; Romanov and Kolesnikov, 2006; Vandenbeuch et al., 2010). Nomura et al., suggest that amiloride-sensitive salt taste is likely mediated by a unique cell type, distinct from Type II or Type III cells, which has also been suggested by others (Chandrashekar et al., 2010; Roebber et al., 2019). It is certainly possible that GAD65Cre drives expression in this unique cell type or progenitors that give rise to this specific taste cell type, which is responsible for sodium-specific salt taste but it is likely not Type I cells.

In summary our data indicate that GAD65Cre activity drives expression of reporter alleles in type II and III taste cells in addition to type I cells, perhaps due to transient expression in taste progenitors. Thus, even though the majority of GAD65Cre/tdTomato+ cells appear to be Type I cells, results using this mouse line to study Type I cell function specifically must be interpreted cautiously

## CONFLICT OF INTERESTS

The authors declare no conflict of interest

## FUNDING

Supported by NIH/NIDCD grant (R01DC017679) to SCK

## ACKNOWLEDGEMENTS

The authors want to thank Drs. Thomas Finger and Linda Barlow for helpful discussions during the course of this study.

## Notes

### Competing Interest Statement

The authors have declared no competing interest.

